# Neuronal Enhancers are Hotspots For DNA Single-Strand Break Repair

**DOI:** 10.1101/2020.12.16.423085

**Authors:** Wei Wu, Sarah E. Hill, William J. Nathan, Jacob Paiano, Kenta Shinoda, Jennifer Colon-Mercado, Elsa Callen, Raffaella de Pace, Dongpeng Wang, Han-Yu Shih, Steve Coon, Maia Parsadanian, Hana Hanzlikova, Peter J. McHugh, Andres Canela, Keith W. Caldecott, Michael E. Ward, André Nussenzweig

## Abstract

Genome stability is essential for all cell types. However, defects in DNA repair frequently lead to neurodevelopmental and neurodegenerative diseases, underscoring the particular importance of DNA repair in long-lived post-mitotic neurons. The neuronal genome is subjected to a constant barrage of endogenous DNA damage due to high levels of oxidative metabolism in the central nervous system. Surprisingly, we know little about the identity of the lesion(s) that accumulate in neurons and whether they accrue throughout the genome or at specific loci. Here, we show that neurons, but not other post-mitotic cells, accumulate unexpectedly high numbers of DNA single-strand breaks (SSBs) at specific sites within the genome. These recurrent SSBs are found within enhancers, and trigger DNA repair through recruitment and activation of poly(ADP-ribose) polymerase-1 (PARP1) and XRCC1, the central SSB repair scaffold protein. Notably, deficiencies in PARP1, XRCC1, or DNA polymerase β elevate the localized incorporation of nucleotides, suggesting that the ongoing DNA synthesis at neuronal enhancers involves both short-patch and long-patch SSB repair processes. These data reveal unexpected levels of localized and continuous DNA single-strand breakage in neurons, suggesting an explanation for the neurodegenerative phenotypes that occur in patients with defective SSB repair.

---

Long-lived neurons are thought to be particularly prone to a progressive accumulation of endogenous DNA lesions. Recent studies have shown that single human neurons harbor large numbers of somatic muations, and that the burden of mutations in neurons substantially increases during aging^1,2^. Neuronal somatic mutations are especially abundant in patients with familial neurodegenerative diseases associated with dysfunctional DNA repair genes^1,3^, as well as neurodevelopmental diseases such as autism and spectrum disorder (ASD)^4^. Despite a wealth of evidence supporting the importance of neuronal DNA damage in neurological diseases, the precise identity and sources of DNA damage that shape the mutational landscape of the neuronal genome remain unclear.

Several different types of DNA damage have been reported to accumulate in neurons including base modifications, single- and double-strand breaks and interstrand crosslinks^3,5^. While defects in all major DNA repair pathways have been implicated in neurological diseases, single-strand break (SSB) repair appears to be particularly important in neurons, because defects in this pathway leads almost exclusively to neuronal dysfunction and degeneration^3,5^. Importantly, DNA excision and DNA strand break repair pathways are associated with unscheduled DNA synthesis, an obligatory and characteristic step of DNA repair known as gap filling. During gap filling, excised or missing nucleotides are replaced, usually using the undamaged strand as a template^6^. Importantly, if gap filling involves the incorporation of sufficient number of nucleotides, DNA repair synthesis can be a robust measure of the extent and sites of endogenous DNA damage^7^.

## Regions of the neuronal genome are associated with ongoing DNA synthesis

Using alkaline comet assays, others previously observed endogenous DNA damage in cultured rat neurons^8^. We developed a new method to measure sites of DNA repair synthesis and map their genomic locations by sequencing (synthesis-associated with repair sequencing; SAR-seq). We labeled post-mitotic iPSC-derived glutamatergic neurons (i^3^Neurons^9,10^) with a thymidine analog EdU for 18 hours, biotinylated the labeled DNA by click chemistry, sonicated it to 150-200 bp, and then isolated the labeled DNA for high-throughput sequencing (**Fig. 1a)**. Surprisingly, we identified >55,000 peaks of DNA synthesis at recurrent genomic locations in neurons, which were highly reproducible between different experiments (**Fig. 1b, Extended Data Fig. 1a, b**). Peaks were not caused by DNA synthesis during S phase^11,12^, because the neurons are post-mitotic and the SAR-seq peaks were unaffected by inhibitors of the replicative DNA polymerases DNA polymerase *α* (PolA), Pol δ, and Pol ϵ (**Extended Data Fig. 1c, d)**. As expected, neuronal DNA synthesis was largely prevented by hydroxyurea, which reduces the availability of deoxyribonucleotides (**Extended Data Fig. 1c, d)**.

**Fig 1:**
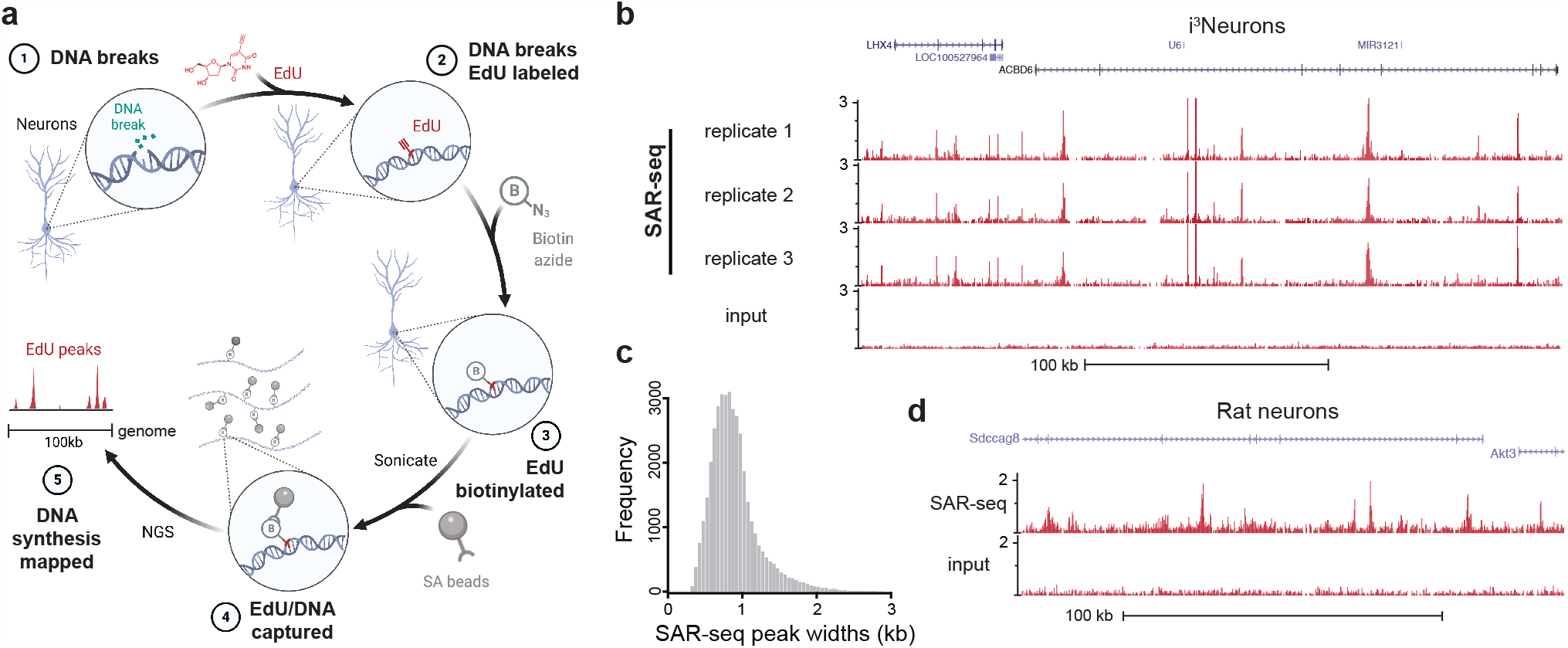
Discrete loci in the neuronal genome are associated with ongoing DNA synthesis. **a)** Schematic of SAR-seq (DNA synthesis associated with repair sequencing) methodology. A population of neurons grown in culture (*1*) are incubated with EdU to label sites of DNA repair synthesis (*2*). Genomic EdU is tagged with biotin via click chemistry (*3*), sheared by sonication and captured with streptavidin beads (*4*). Enriched DNA sequences are PCR-amplified and subjected to next-generation sequencing (*5*). **b)** Genome browser screenshot displaying SAR-seq profiles as normalized read density (reads per million, RPM) for human iPSC-derived neurons (i^3^Neurons). Three independent biological replicates are shown as well as input DNA sequenced in parallel. **c)** Histogram of individual SAR-seq peak widths, with an average width of 1,006 bp. **d)** Genome browser screenshot of SAR-seq performed on rat primary neurons. The culture was simultaneously treated with 5μM aphidicolin to block DNA replication in S phase glial cells.

The tracts of DNA synthesis ranged from about 200-2,000 bp in width (averaging 1,006 bp) (**Fig. 1c**), most likely reflecting multiple clustered sites of DNA repair. The most prominent SAR peaks in the post-mitotic neurons were detectable by pulse labeling with EdU for just 1 hour, and EdU incorporation approached saturation after labeling for 18 hours (**Extended Data Fig. 2a)**. On average, we estimate that any given tract was labeled to 89% capacity within ∼10 hours of EdU labeling (**Extended Data Fig. 2b**).

**Fig 2:**
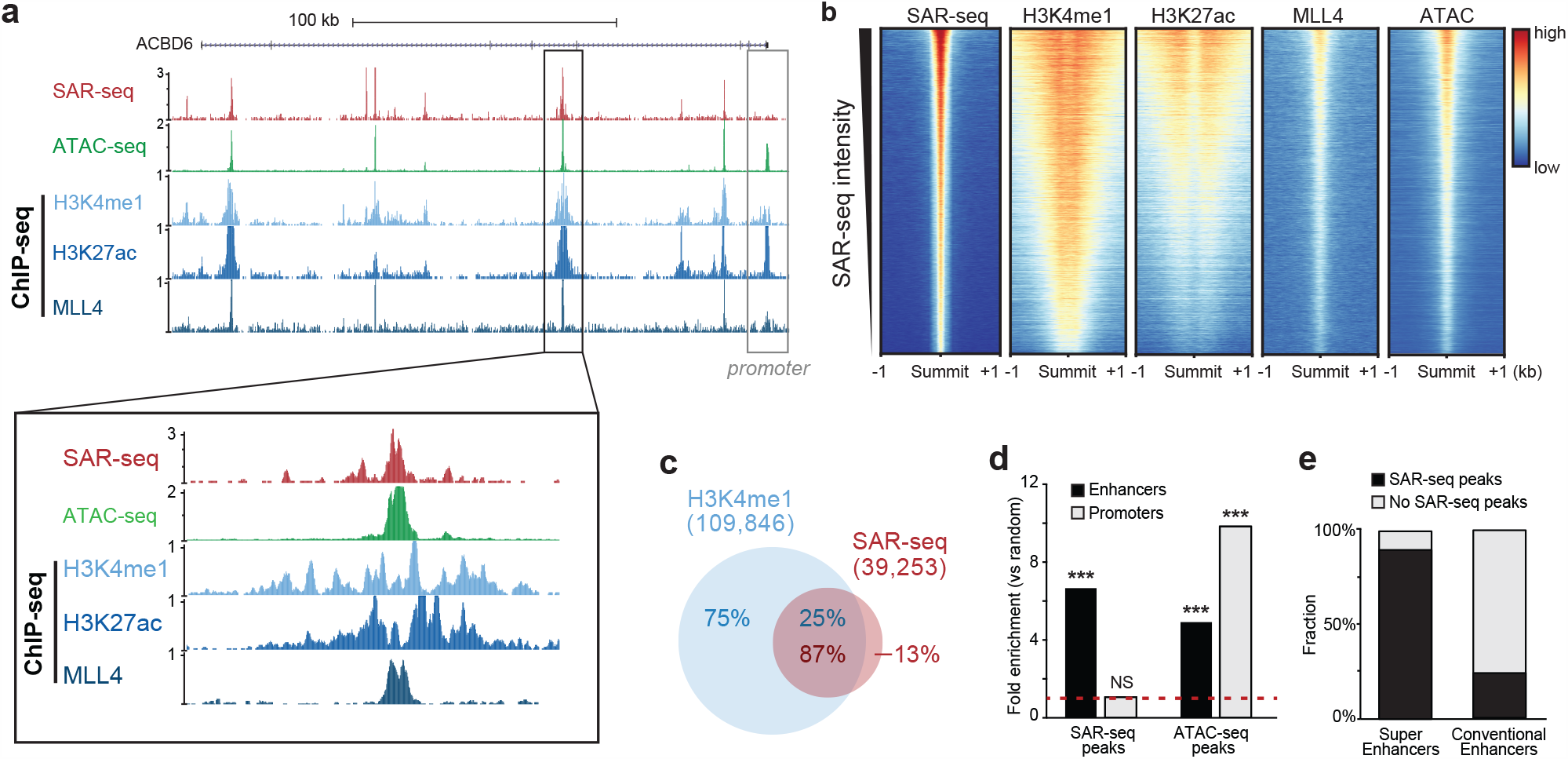
SAR-seq peaks occur within nucleosome-free regions of enhancers. **a)** Genome browser screenshots displaying SAR-seq (red), ATAC-seq (green), and H3K4me1 (light blue), H3K27ac (steel blue), and MLL4 ChIP-seq (indigo) in i^3^Neurons. Zoomed-in screenshot of the highlighted region is shown below. ATAC-seq and MLL4 peaks align directly with the SAR-seq peak, while H3K4me1 and H3K27ac peaks flank the SAR-seq site. Region highlighted in black in top panel shows the absence of SAR-seq signal at the promoter (see also quantitation in Fig. **2d** and Extended Data Figs. **2e** and **3d** below). **b)** Heatmaps of SAR-seq, H3K4me1, H3K27ac, MLL4 ChIP-seq and ATAC-seq signal ±1kb around SAR-seq peak summits in i^3^Neurons, ordered by SAR-seq intensity. SAR-seq peaks are located within the nucleosome-free region of H3K4me1 and H3K27ac ChIP-seq peaks, and colocalize with MLL4 ChIP-seq and ATAC-seq regions. **c)** Venn diagram showing the overlap between H3K4me1 and SAR-seq peaks in i^3^Neurons. n= 1,000 random datasets were generated to test the significance of overlap (one-sided Fisher’s Exact test (p<2.2e-16)). **d)** Graph showing the fold enrichment of SAR-seq and ATAC-seq peaks located at enhancers (black) and promoters (grey) compared to 1000 sets of randomly shuffled regions of the same sizes and chromosome distributions, respectively. (Fisher’s Exact test ***p<2.2e-16; NS, p=0.07)). **e)** Graph showing the fraction of super-enhancers (left) overlapping with SAR-seq peaks compared to conventional enhancers (right). 1,385 super-enhancers were defined by H3K27ac ChIP-seq intensity in i^3^Neurons (see **Extended Data Fig. 4d**).

Importantly, when iPSCs were differentiated into skeletal muscle cells rather than neurons, we did not detect incorporation of EdU (**Extended Data Fig. 2c**). Similarly, we failed to detect EdU incorporation in G0-arrested pre-B cells, although we could detect EdU incorporation in the pre-B cells at site-specific DNA double strand breaks (DSBs) (**Extended Data Fig. 2d)**. To rule out the possibility that the SAR-seq peaks were an artifact of iPSC differentiation, we also labelled *bona fide* rat neurons with EdU. Similar to i^3^Neurons, we detected robust peaks of EdU incorporation at specific sites in the rat neurons (**Fig. 1d**). Thus, the high frequency of recurrent DNA synthesis appears to be a specific feature of post-mitotic neurons.

## The recurrent sites of neuronal DNA synthesis are located in cis-acting enhancers

Neuronal SAR-seq peaks were highly enriched in intragenic regions (**Extended Data Fig. 2e**) and associated with expressed genes (**Extended Data Fig. 2f**), suggesting the involvement of transcription. However, the signal intensity did not correlate with transcript levels as measured by RNA-seq (**Extended Data Fig. 2f**). Moreover, the sites of EdU incorporation were not associated with strand-specificity, because the EdU was incorporated uniformly in both transcribed and non-transcribed strands, resulting in symmetrical strand-specific SAR-seq profiles (**Extended Data Fig. 2g**).

We next searched for specific DNA motifs using the strongest 5,000 SAR-seq peaks. More than 50% of the sites harbored a motif similar to the ONECUT family of transcription factors, and the ONECUT1 motif was centered at SAR-seq peaks (**Extended Data Fig. 3a**). Since ONECUT1 has been implicated in promoting genomic accessibility in neurons^13^, we compared SAR-seq peaks with accessible regions using ATAC-seq. We found that 57% of the SAR-seq regions coincided with ATAC-seq peaks (**Extended Data Fig. 3b**) and the width of the SAR-seq peaks correlated with the width of the ATAC-seq peaks, suggesting that open chromatin structure influences the extent of DNA synthesis (**Extended Data Fig. 3c and Fig. 1c**).

**Fig 3:**
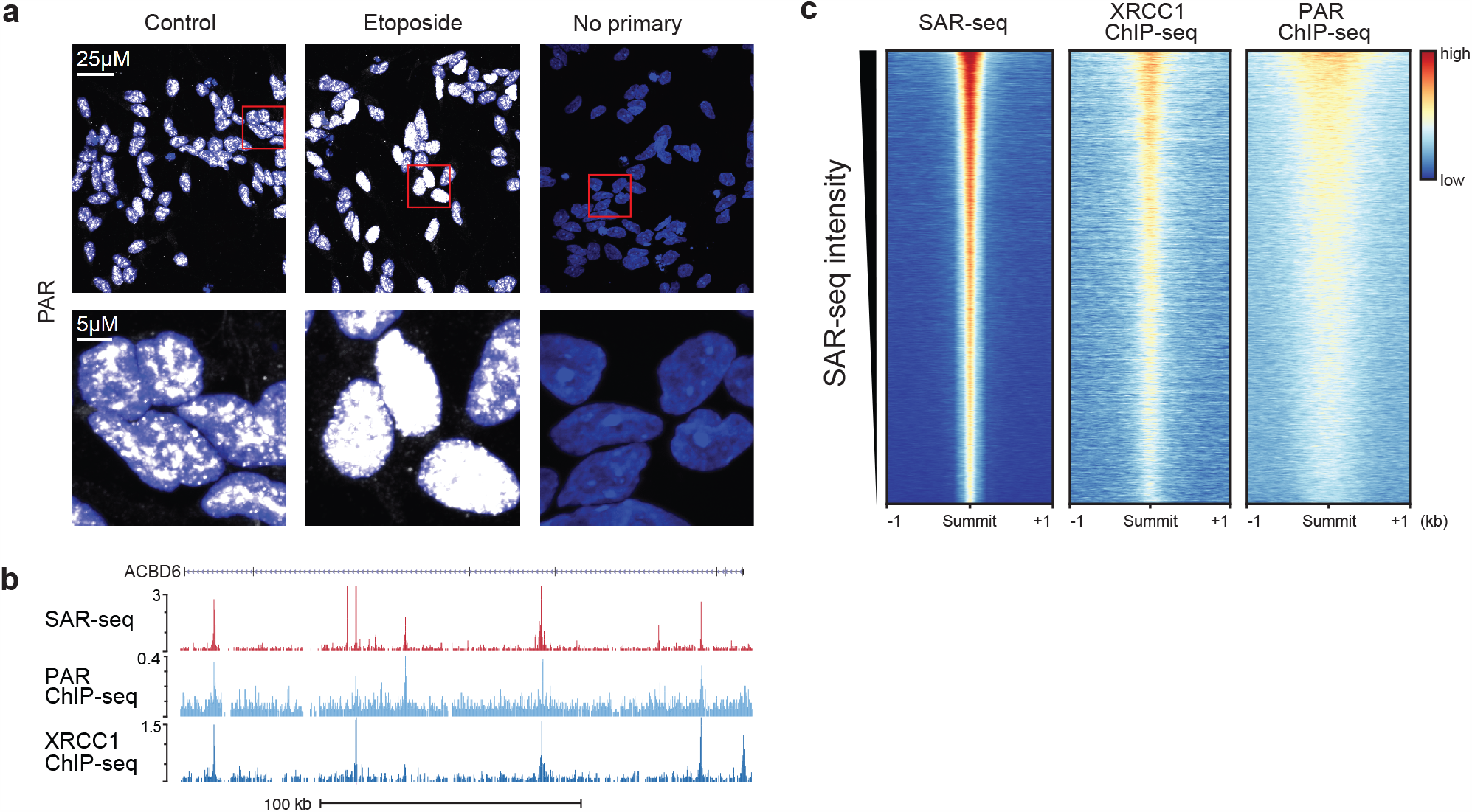
PARP activation and XRCC1 recruitment at neuronal sites of DNA synthesis. **a)** Representative images of anti-PAR immunofluorescence staining in i^3^Neurons. As a positive control for DNA damage and increased PARylation, cells were treated with 50uM etoposide (ETO) for 18 hr. Primary anti-PAR antibody without secondary antibody was used as a negative control. **b)** Genome browser screenshots displaying SAR-seq (red), PAR (light blue) and XRCC1 ChIP-seq (steel blue) signal in i^3^Neurons. Cells were incubated with PARGi 20 minutes prior to fixation for PAR ChIP-seq. **c)** Heatmaps of XRCC1 and PAR ChIP-seq signal ±1kb surrounding SAR-seq peak summits in i^3^Neurons, ordered by SAR-seq intensity.

Although the neuronal sites of recurrent DNA synthesis are located predominantly in open chromatin, SAR-seq peaks were not enriched at promoters like ATAC-seq peaks (**Fig. 2a**,**d and Extended Data Fig. 3d**). Indeed, promoters occupied by RNA polymerase II (POL II), as determined by ChIP-seq, exhibited only modest levels of DNA synthesis (**Extended Data Fig. 3e**). In contrast to promoters, however, we detected a strong positive correlation between the location of DNA synthesis and neuronal enhancers, as measured by ChIP-seq for H3K4me1, H3K27ac and MLL4, the major mammalian histone H3K4 mono-methyltransferase^14^ (**Fig. 2a-c**). Indeed, SAR-seq peaks were centered on the nucleosome-free region occupied by MLL4 (**Fig. 2a, b**). Altogether, 87% of SAR-seq peaks overlapped with H3K4me1 peaks and were significantly enriched at enhancers compared to random regions (**Fig. 2c, d**), indicating enhancers are hotspots of neuronal DNA synthesis. Consistent with these results in i^3^Neurons, SAR peaks in primary rat neurons also overlapped and correlated with enhancers **(Extended Data Fig 4a)**.

**Fig 4:**
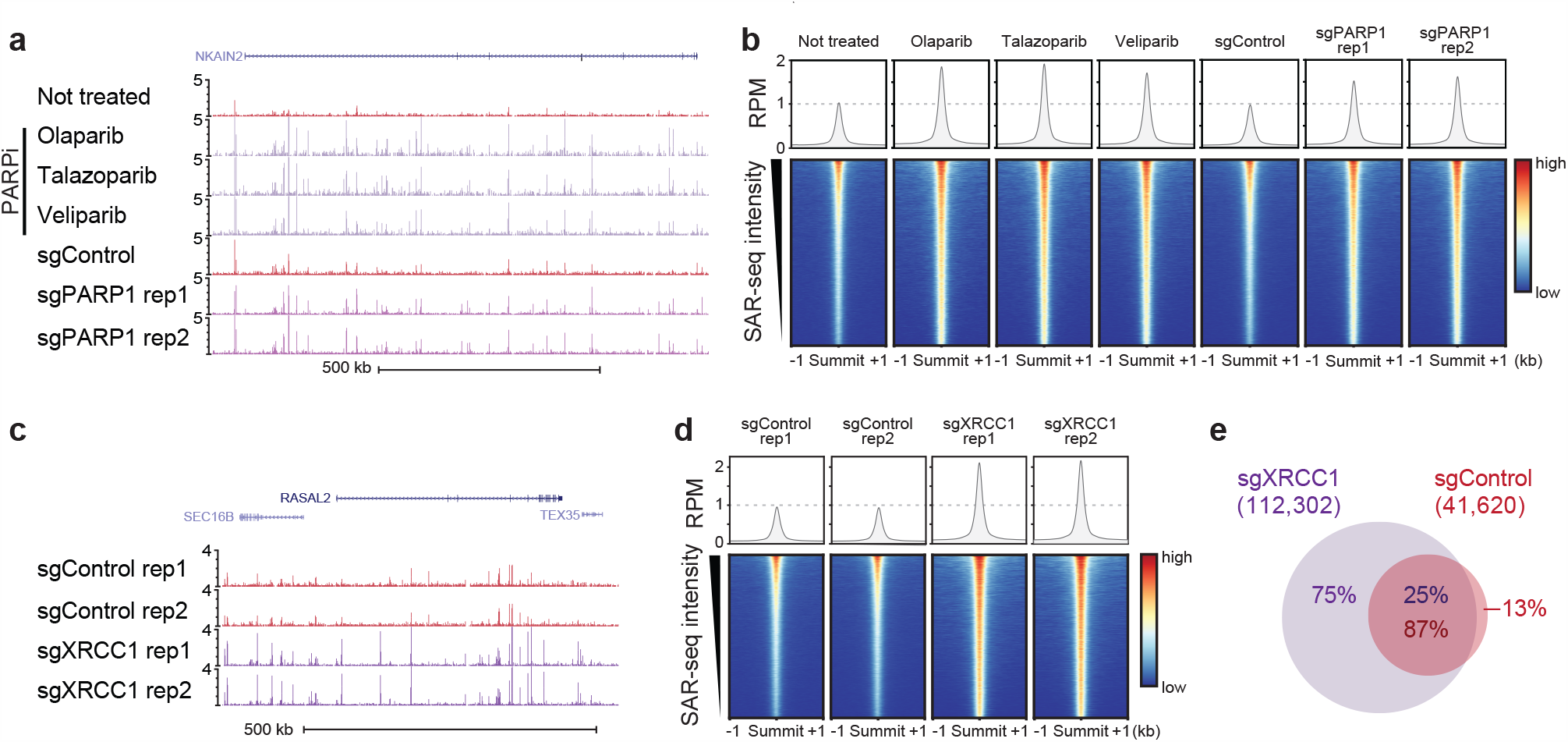
PARP or XRRC1 deficiency increases SAR-seq peak intensity. **a)** Genome browser screenshots displaying SAR-seq profiles in i^3^Neurons treated with PARP inhibitors olaparib, talazoparib, veliparib, or CRISPRi-mediated knockdown with a control non-targeting sgRNA (sgControl) or an sgRNA targeting PARP1 (sgPARP1) in duplicates. **b)** Heatmaps of SAR-seq intensities ±1kb surrounding SAR-seq peak summits for i^3^Neurons treated with PARP inhibitors and in neurons containing CRISPRi-mediated PARP1 knockdown or non-targeting sgRNA control. Aggregate plot of SAR-seq intensity is shown in the top panel. **c)** Genome browser screenshots of SAR-seq profiles in i^3^Neurons expressing CRISPRi non-targeting sgRNAs or targeting XRCC1 (sgXRCC1), in duplicate. **d)** Heatmaps of SAR-seq intensities ±1kb surrounding SAR-seq peak summits for i^3^Neurons expressing non-targeting sgRNA or targeting XRCC1. Aggregate plot of SAR-seq intensity is shown in the top panel. **e)** Venn diagrams showing the overlap of SAR-seq peaks between control (red) and XRCC1 (purple). n = 1,000 random datasets were generated to test significance of overlap (one-sided Fisher’s Exact test (p<2.2e-16)).

MLL4 has been reported to colocalize with lineage determining transcription factors^14^. To determine whether the enhancers with SAR-seq peaks are specific to those that are active in differentiated neurons, we compared sites of H3K4me1 in neurons with those in the iPSCs from which they were derived. Only 2% of the SAR-seq peaks overlapped with iPSCs-specific H3K4me1peaks, while most of the SAR-seq peaks overlapped with either neuron-specific or H3K4me1 sites that were shared between neurons and iPSCs (**Extended Data Fig. 4b**). Thus, based on H3K4me1 localization, SAR is associated with enhancers that are active in differentiated neurons.

We then performed Gene Ontology (GO) analysis to determine biological processes associated with the genes containing SAR-seq peaks. Many of the top enriched GO terms related to neuronal migration, development, axon formation, and synapse assembly, reflecting nervous system function (**Extended Data Fig. 4c**). Super-enhancers (SE) are a large collection of enhancers that drive transcription of genes involved in cell identity. We found approximately 1,300 SE in i^3^Neurons as determined by H3K27ac ChIP-seq, 90% of which exhibited SAR-seq peaks, whereas less than 25% of conventional enhancers were enriched in SAR (**Fig. 2e, Extended Data Fig. 4d**). Collectively, these data identify super-enhancers and genes associated with neuronal function as hotpsots of recurrent DNA synthesis.

## PARP activation at neuronal sites of DNA synthesis

Next, we addressed the source of the DNA synthesis in post-mitotic neurons. Given the close association of unrepaired DNA breaks with neurodegeneration, we wondered if the sites of EdU incorporation might reflect sites of DNA break repair. To test this, we measured the activity of poly(ADP-ribose) polymerase (PARP) enzymes at the sites of DNA synthesis. PARP1 and PARP2 are activated in response to various types of DNA breakage, including SSBs, DSBs, and single-strand gaps^15,16^. PARP activity signals the presence of these lesions by modifying proteins with poly(ADP-ribose) (PAR), after which PAR is rapidly removed from proteins by poly(ADP-ribose) glycohydrolase (PARG)^15,16^. When we monitored ADP-ribosylation in individual neurons using an anti-pan-ADP ribose-binding reagent,^17^ we observed robust focal staining in neuronal nuclei (**Fig. 3a**). To determine whether the poly(ADP-ribose) was located at sites of neuronal DNA synthesis we employed PAR ChIP-seq, which we demonstrated can detect ADP-ribosylation at site-specific DSBs in pre-B cells (**Extended Data Fig. 5a**). Strikingly, when we used this method in i^3^Neurons, we found that sites of PAR synthesis co-localized with sites of DNA synthesis (**Fig. 3b, c)**. These findings strongly suggest that the recurrent sites of DNA synthesis at neuronal enhancers are associated with sites of DNA break repair.

**Fig 5:**
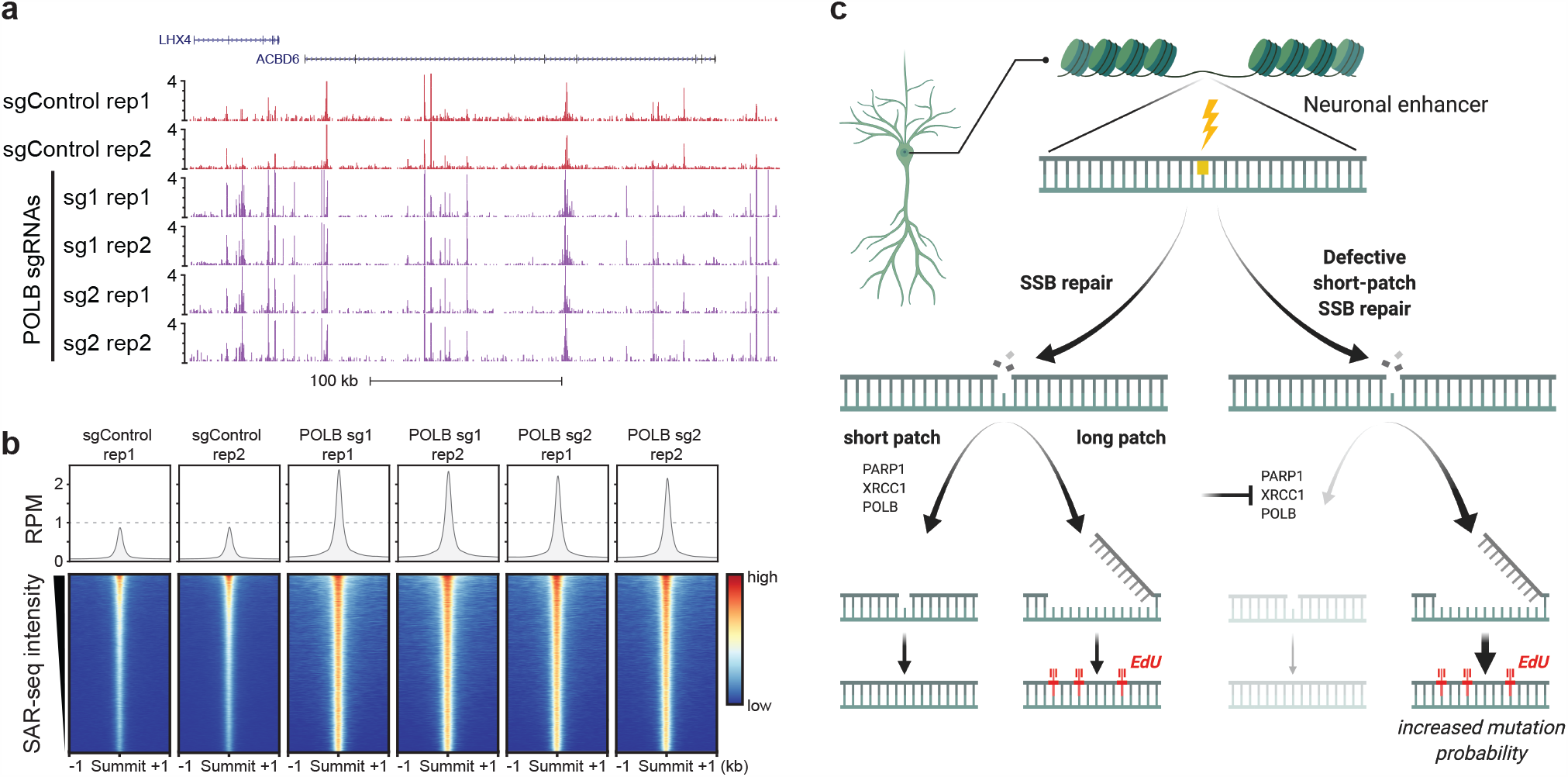
Deficiency in POLβ elevates the localized incorporation of EdU. **a)** Genome browser screenshots displaying SAR-seq profiles in i^3^Neurons expressing CRISPRi-mediated knockdown of POL*β* (sgPOLB, two guides are shown) or a control non-targeting sgRNA (sgControl), in duplicates. **b)** Heatmaps of SAR-seq intensities ±1kb surrounding SAR-seq peak summits for i^3^Neurons expressing CRISPRi-mediated knockdown of POL*β* or a control non-targeting sgRNA. Aggregate plot of SAR-seq intensity is shown in the top panel. **c)** Model suggesting that PARP1, XRCC1, and POL*β* influence the choice between SP- and LP-SSB repair. Loss of these proteins results in patches of increased DNA synthesis arising from LP-SSB repair, which may increase mutation frequency.

## Sites of neuronal DNA synthesis are sites of DNA single-strand break repair

Neuronal activity has been reported to result in DSBs^18,19^. Moreover, topoisomerase 2 (TOP2) is associated with neuronal activity. TOP2-induced DSBs can promote the expression of early response genes^18^, associating these DNA breaks with regions of transcriptional activity. However, while we found that treatment of i^3^Neurons with etoposide to trigger TOP2-induced DSBs resulted in DNA synthesis within gene bodies, most of the sites of etoposide-induced DNA synthesis were distinct from those detected in untreated neurons (**Extended Data Fig. 5b, c**). In addition, we did not detect DSBs in unchallenged i^3^Neurons as measured by either *γ*-H2AX/53BP1 immunostaining (**Extended Data Fig. 5d**) or by END-seq (**Extended Data Fig. 5e**), which maps DSBs genome-wide at base-pair resolution^20^. Thus, the sites of DNA synthesis in neuronal enhancers are independent of DSBs.

Importantly, PARP1 and/or PARP2 are also activated at SSBs, which subsequently recruits the XRCC1 protein complexes necessary for the repair of DNA single-strand gaps and breaks^21,22^. We therefore examined the genomic localization of XRCC1 by ChIP-seq. Strikingly and similar to PAR, XRCC1 also co-localized with SAR-peaks, and the intensity of XRCC1 binding was correlated with the intensity of EdU incorporation in both human i^3^Neurons and primary rat neurons (**Fig. 3b, c; Extended Data Fig. 5f**). Collectively, these data indicate that the sites of DNA synthesis at neuronal enhancers are sites of PARP activation and XRCC1-associated SSB repair.

## DNA synthesis at neuronal enhancers involves both short-patch and long-patch SSB repair

PARP1 and XRCC1 promote the repair of a wide spectrum of SSBs^23^. We therefore examined the impact of inhibiting and/or depleting these proteins on neuronal DNA synthesis. Intriguingly, we observed a reproducible increase in SAR when neurons were treated with three independent inhibitors of PARP during the EdU incubation, or if PARP1 was depleted using CRISPR interference (CRISPRi)^24^ (**Fig. 4a, b, Extended Data Fig. 6a**). Moreover, depletion of XRCC1 similarly led to a prominent increase in neuronal SAR (**Fig. 4c-e; Extended Data Fig. 6b**). These data suggest that if PARP1- and XRCC1-dependent repair is impeded, the amount of EdU incorporation that is associated with SSB repair is increased. The simplest explanation for these data is that in the absence of XRCC1-dependent short-patch repair, long-patch repair is triggered instead **(Fig. 5c)**. During short-patch SSB repair a single nucleotide is replaced at the site of the break^23,25^, typically by DNA polymerase *β* (POL*β*) which interacts directly with XRCC1^5,21^ **(Fig. 5c)**. In contrast, during long-patch repair, alternative DNA polymerases can achieve gap filling, at the expense of increased patch sizes for DNA synthesis. Indeed, we found that depletion of POL*β* by two independent CRISPRi sgRNAs resulted in a dramatic increase in DNA synthesis at neuronal enhancers (**Fig. 5a, b; Extended Data Fig. 6c, d**). Thus, the increased DNA repair synthesis in PARP1/XRCC1/POL*β*-depleted neurons is consistent with increased activity of long-patch SSB repair.

## Conclusions

Our study reveals that human post-mitotic neurons are subject to a profound and unexpected degree of localized and recurrent DNA synthesis, which is associated with ongoing sites of SSB repair at neuronal enhancers. The scaffold protein XRCC1 is of particular importance during SSB repair because it interacts directly with many of the components of short-patch repair including POL*β* and LIG3^21^. Interestingly, patients with hereditary mutations in XRCC1 exhibit progressive neurodegeneration, suggesting that intact SP-SSB repair is necessary for long-term neuronal viability^26,27^. One possible explanation for this is that the elevated PARP1 activity that results from loss of XRCC1-dependent SSB repair triggers neuronal dydsfunction and/or ultimately cell death^21^. In addition, our current data raise the possibility that an increased dependency on long-patch DNA repair synthesis in XRCC1-defective cells increases mutational burden in long-lived neurons (**Fig. 5c**). Age-dependent accrual of mutations at neuronal enhancers through hyperactive LP-SSB repair could subsequently lead to aberrant transcription, eventually resulting in neurodegeneration. This notion is consistent with the recent finding that somatic mutations measured in single human neurons are enriched at highly-accessible genomic regions including brain-specific enhancers and genes regulating neuronal function^1,2,4^. Although the mechanism(s) for such mutations were not defined, three distinct somatic mutation signatures were identified: a clock-like signature of aging, a brain region-specific signature, and a mutation class most closely associated with oxidative DNA damage^1,2^.

It is unexpected that sites of endogenous SSB repair would be enriched at neuronal enhancers rather than being randomly distributed. Enhancers could be especially vulnerable to single-strand damage due to their increased mobility upon transcriptional activation^28^, or because negatively-charged products of oxidative metabolism might preferentially interact with phase-separated enhancer condensates^29^. Intriguingly, active DNA demethylation of cytosine at CpG sites occurs preferentially at enhancers^30^, is 10-fold more active in post-mitotic neurons than peripheral cell types ^31^, and generates SSBs that are intermediates of XRCC1-associated BER^30,32^. Indeed, we have found that CpG dinucleotides are highly enriched at SAR sites (**Extended Data Fig. 7**). While the short patch-BER pathway that replaces methylated cytosines would not generate EdU incorporation, DNA synthesis surrounding this site by long patch-BER might be a frequent outcome.

In summary, we describe a new method that enables genome-wide mapping of sites of DNA repair in post-mitotic cells. Our findings identify enhancers as hotspots of endogenous DNA single-strand break repair in human post-mitotic neurons, and may explain why this important DNA repair process is crucial for maintaining the functionality and viability during neuronal aging.

Note: During the preparation of this manuscript, we became aware of the closely related work of D. Ried *et al*., which demonstrated recurrent DNA repair sites in embryonic stem cell-derived neurons. doi: https://doi.org/10.1101/2020.03.25.008490

## Acknowledgments

We thank Michael Kruhlak, Yilun Sun, Yves Pommier, and Kai Ge for helpful discussions and reagents; KWC is supported by Programme Grants from the UK Medical Research Council (MR/P010121/1), Cancer Research-UK (C6563/A7322), and by ERC Advanced Investigator (SIDSCA 694996) and Royal Society Wolfson Research Merit Awards; The M.W. laboratory is supported by the NINDS Intramural Research Program. S.E.H. received funding from the Brightfocus Foundation. The A.N. laboratory is supported by the Intramural Research Program of the NIH, an Ellison Medical Foundation Senior Scholar in Aging Award (AG-SS-2633-11), the Department of Defense Awards (W81XWH-16-1-599 and W81XWH-19-1-0652), the Alex’s Lemonade Stand Foundation Award, and an NIH Intramural FLEX Award.

## Author Contributions

W.W., S.E.H., W.J.N., J.P., K.W.C., M.E.W., and A.N. designed the study. S.E.H., W.J.N., J.P., performed most of the experiments with assistance from K.S., J.C-M., E.C., R.P., D.W., H.-Y., S.C., M.P. on certain experiments. A.C. developed SAR-seq in the A.N. lab, W.W. designed bioinformatics pipelines and performed data analysis, and designed the figures. H.H., P.J.M and A.C provided insights. K.W.C., M.E.W., and A.N. analyzed and interpreted data and wrote the paper. W.W., S.E.H., W.J.N., J.P., helped edit the paper. M.E.W. and A.N. supervised the study.

---

**Extended Data Fig 1:**
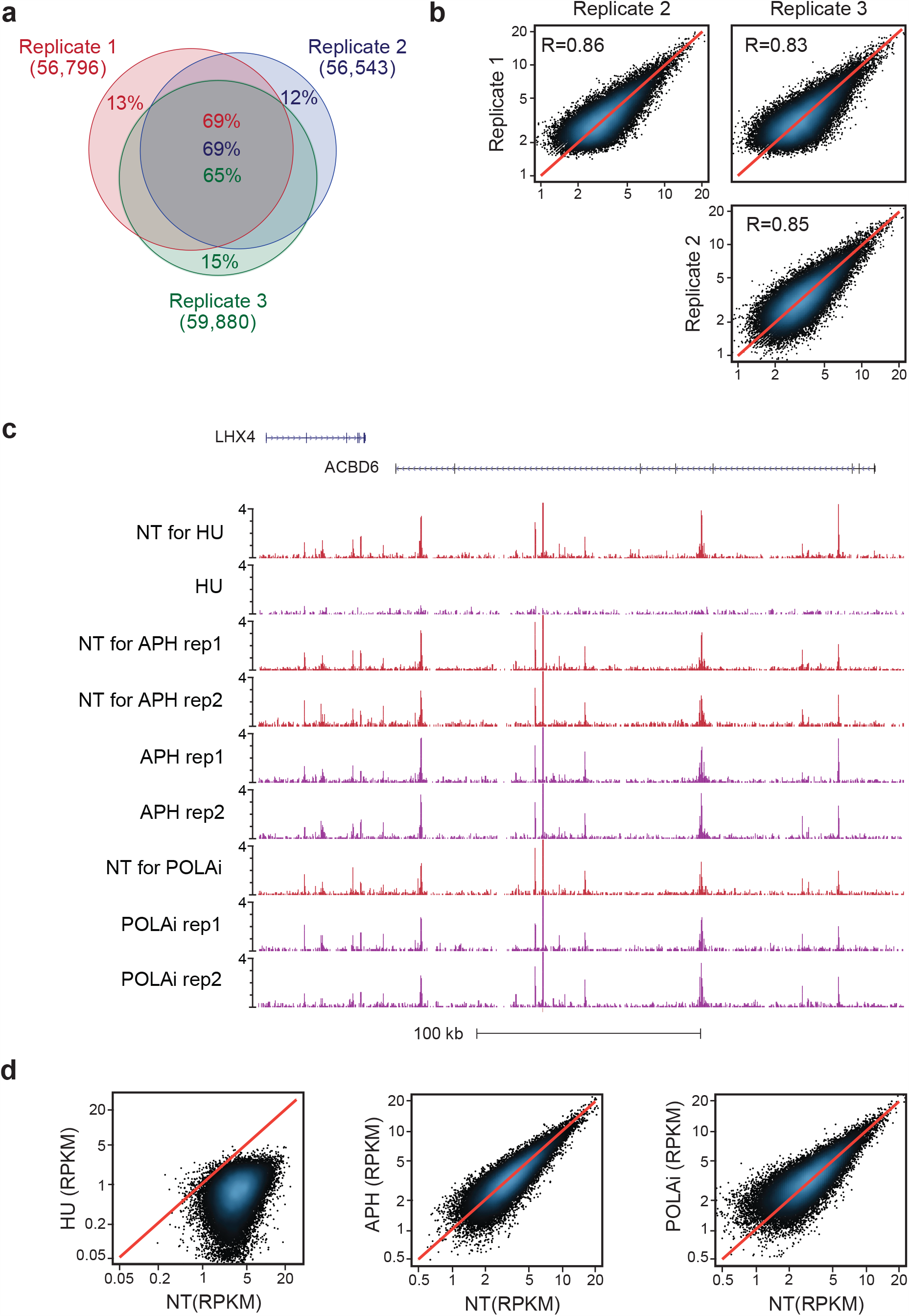
Mapping regions of unscheduled DNA synthesis in neurons. **a)** Venn diagram showing the overlap of SAR-seq peaks in i^3^Neurons in three independent biological replicates. **b)** Correlations of SAR-seq intensities between three replicates in i^3^Neurons. Pearson correlation coefficient is indicated. **c)** Genome browser screenshot showing SAR-seq in i^3^Neurons treated with hydroxyurea (HU), aphidicolin (APH) or polymerase alpha inhibitor (POLAi). NT, not treated. **d)** Scatterplots showing SAR-seq intensity (SAR-seq reads per kilobase per million mapped reads, RPKM) for HU-(left), and APH (middle)- and POLAi-(right) treated compared to non-treated samples.

**Extended Data Fig 2:**
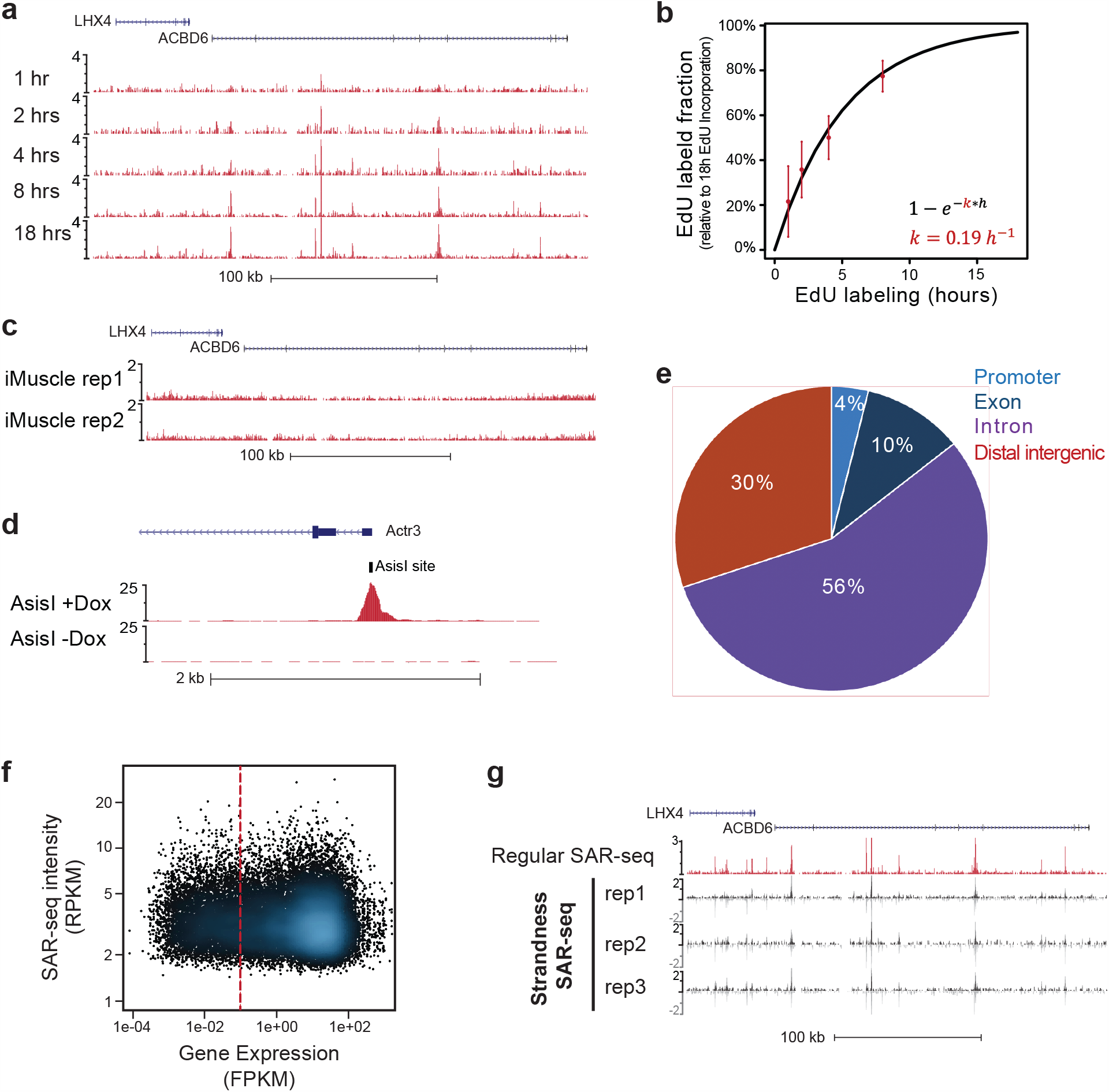
Genomic features of SAR-seq peaks. **a)** Genome browser screenshot showing SAR-seq in i^3^Neurons harvested after 1 hour, 2 hours, 4 hours, 8 hours, or 18 hours of EdU incubation in otherwise non-treated cells. **b)** Graph showing the fraction of EdU labeled in i^3^Neurons (relative to maximum labeling at 18 hours) as a function of time. Red points and error bars represent the relative levels of EdU labeled from experimental data. Black line represents the theoretical model after fitting, with *k* being the rate of EdU labeling. **c)** Genome browser screenshot showing SAR-seq in iMuscle cells for two independent biological replicates. Cells were pretreated with aphidcolin. **d)** Genome browser screenshot displaying SAR-seq peak at a representive AsiS1 restriction enzyme sites (tick mark). AsiS1 expression was induced for 18 hrs (+Dox) vs non-treated (-Dox) in G0-arrested, Abelson virus-transformed murine pre-B cells as described ^20^. **e)** Pie-chart showing distribution of i^3^Neuron SAR-seq peaks at promoters, exons, introns and distal intergenic regions. Promoters are defined as 1kb upstream of transcription start sites and distal intergenic represents promoter-excluded intergenic regions. **f)** Intensity of SAR-seq peaks within intragenic regions compared with transcript levels measured by RNA-seq. 71% of SAR-seq peaks are at expressed genes (FPKM≥0.1, red dashed line). **g)** Genome browser screenshot comparing SAR-seq vs. strand-specific SAR-seq that discriminates which strand is labeled with EdU. Both strands show similar labeling, in triplicate.

**Extended Data Fig 3:**
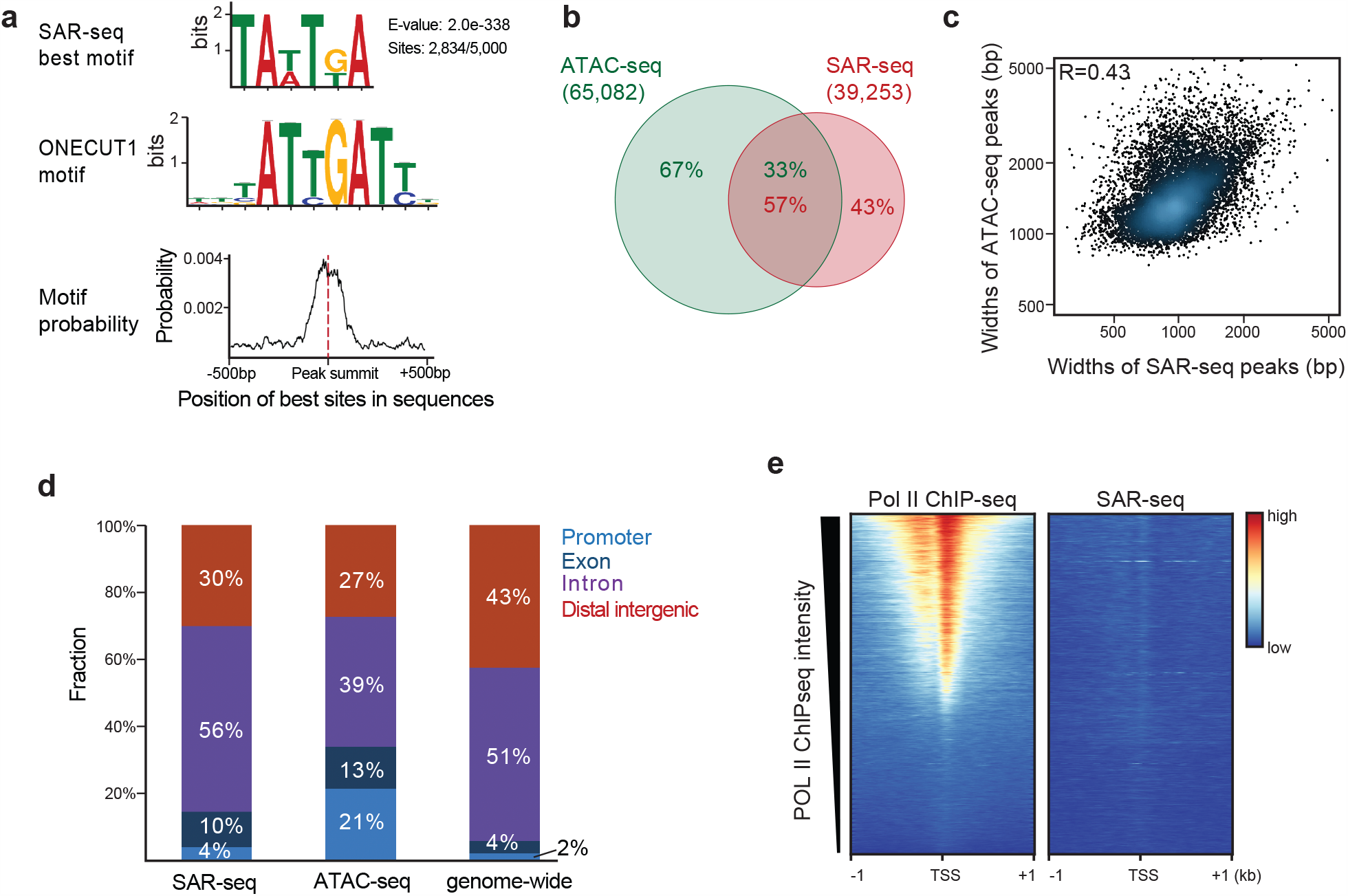
Motif of SAR-seq peaks and comparison with ATAC-seq peaks. **a)** Motif analysis for the sequences within ±500bp surrounding the summit of top 5,000 SAR-seq peaks in i^3^Neurons. Upper panel, the best motif discovered by the MEME suite analysis. 2,834 out of 5,000 sites shared this motif. Middle panel, the motif of the transcription factor ONECUT1 was identified as the best matched motif to SAR-seq motif by the TOMTOM motif comparison tool. Bottom panel, position distribution of the best motif (shown in the upper panel) at ±500bp window of the SAR-seq peak summit. The best motif is centered on the SAR-seq peak summit. **b)** Venn diagram showing overlap between peaks detected in ATAC-seq (green) and SAR-seq (red) in i^3^Neurons. n = 1,000 random datasets were generated to test significance of overlap (one-sided Fisher’s Exact test (p<2.2e-16)). **c)** Scatterplot comparing widths of ATAC-seq peaks and SAR-seq peaks in i^3^Neurons. Pearson correlation coefficient is indicated. **d)** Distribution of SAR-seq and ATAC-seq peaks with respect to different genomic features compared to genome-wide distribution in the hg19 human reference genome. Promoters are defined as 1kb upstream of transcription start sites and distal intergenic represents promoter-excluded intergenic regions. **e)** Heatmaps of RNA Polymerase II ChIP-seq signal and SAR-seq signal within ±1kb surrounding the transcription start site (TSS) in i^3^Neurons, ordered by RNA Polymerase II ChIP-seq intensity.

**Extended Data Fig 4:**
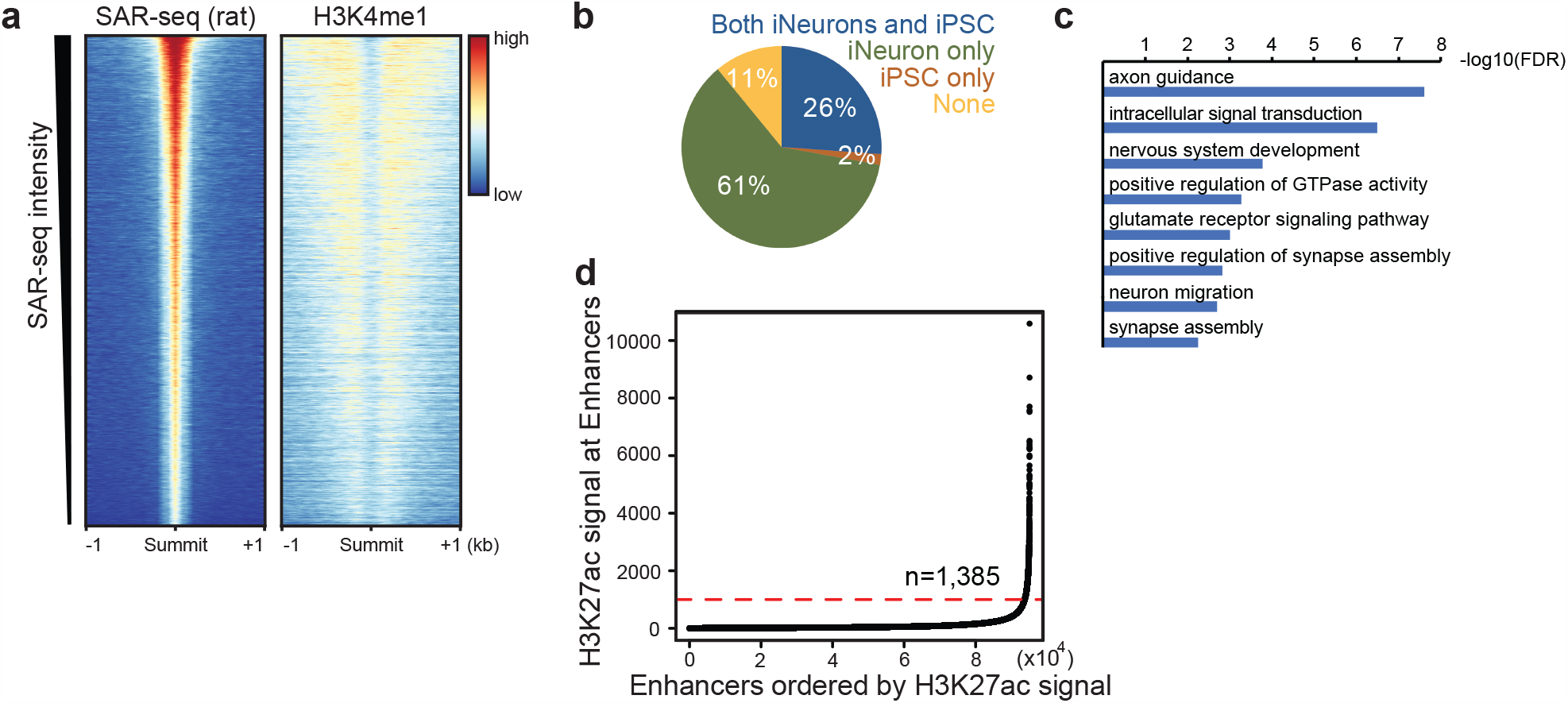
SAR enrichment at primary neuron enhancers and i^3^Neuron super-enhancers. **a)** Heatmaps of SAR-seq and H3K4me1 ChIP-seq signal at ±1kb around the SAR-seq peak summit in embryonic day 18 primary rat cortical neurons ordered by SAR-seq intensity. **b)** Pie-chart showing distribution of i^3^Neuron SAR-seq peaks in iPSC-specific, i^3^Neuron-specific and iPSC- and i^3^Neuron-shared enhancers. **c)** Top biological processes enriched for the genes containing top 2000 SAR-seq peaks determined by Gene Ontology (GO) analysis. **d)** H3K27ac signal at enhancers in i^3^Neurons ranked by H3K27ac ChIP-seq intensity. Red dashed line indicates the inflection point of H3K27ac signal used to determine super enhancers (cutoff: 1000). 1,385 enhancers were defined as super enhancers.

**Extended Data Fig 5:**
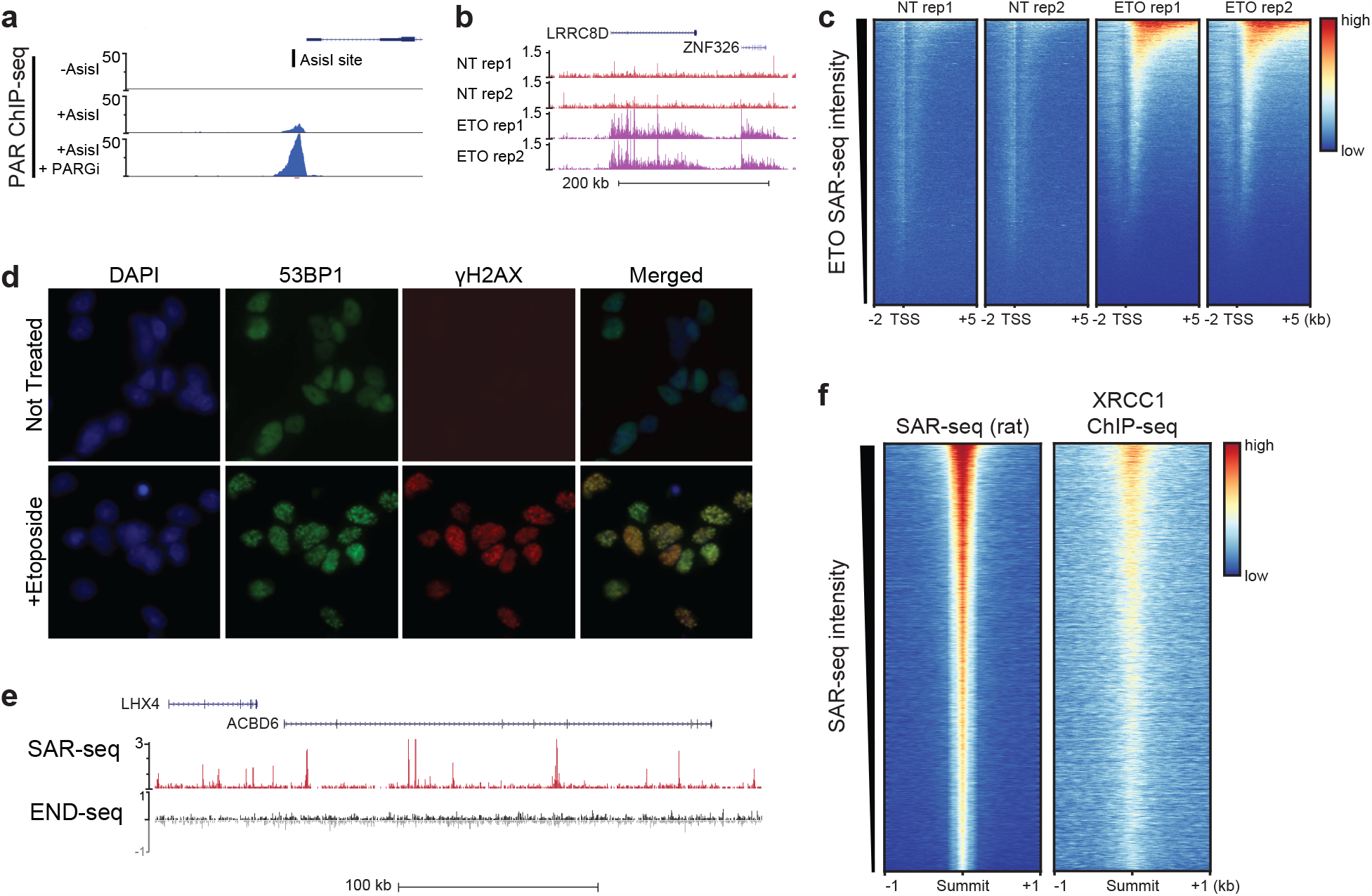
Mapping regions of DNA damage and repair in neurons. **a)** PAR ChIP-seq reads at AsiSI restriction enzyme cut site in Abelson virus-transformed murine pre-B cells. Cells were arrested in G0, and AsiSI double-strand breaks were induced for 18 hr prior to ChIP fixation. PARylation is not observed in non-treated (-AsiSI) cells and is robustly increased after 20 min PARGi treatment prior to fixation (AsiSI + PARGi). **b)** Genome browser examples of SAR-seq in either control (NT) or 18 hr 50uM etoposide (ETO) treated neurons, in duplicate. **c)** Heatmaps for SAR-seq in either control (NT) or in 18 hrs 50uM etoposide-(ETO) treated i^3^Neurons at −2kb to +5kb of the transcription start sites (TSS) ordered by ETO SAR-seq signal. **d)** Immunofluorescence staining of DSB markers *γ*-H2AX and 53BP1 in non-treated or 1 hour etoposide-treated i^3^Neurons **e)** SAR-seq vs END-seq in non-treated i^3^Neurons. END-seq, a method to specifically detect double-strand breaks, shows no enriched signal over background at SAR-seq peaks. END-seq reads are separated by plus and minus DNA strands (black and grey tracks, respectively). **f)** Heatmaps of SAR-seq and XRCC1 ChIP-seq in cultured primary rat neurons, centered on SAR-seq peak summits and ordered by SAR-seq intensity.

**Extended Data Fig 6:**
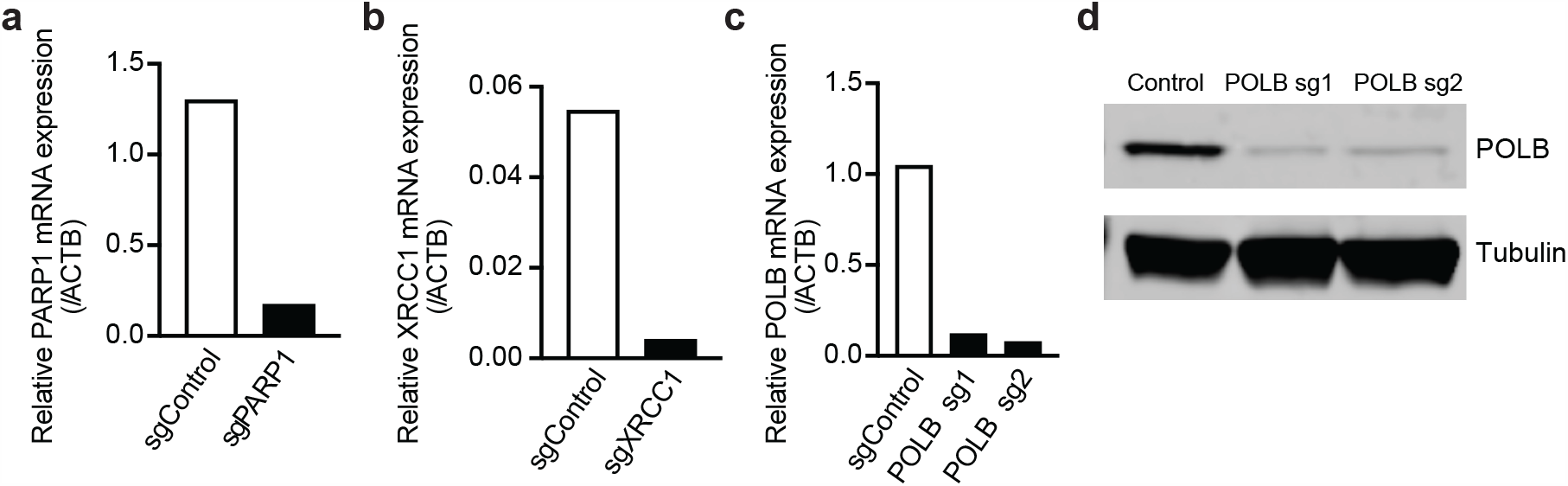
Confirmation of PARP1, XRCC1, and POLb knockdowns in i^3^Neurons. (**a-c**) Quantitative RT-PCR analysis of PARP1 (**a**), XRCC1 (**b**), and POL*β* (**c**) transcript levels in i^3^Neurons after CRISPRi knockdown, cultured in parallel with samples used for SAR-seq. **d**) Western blot of POL*β* protein after CRISPRi knockdown.

**Extended Data Fig 7:**
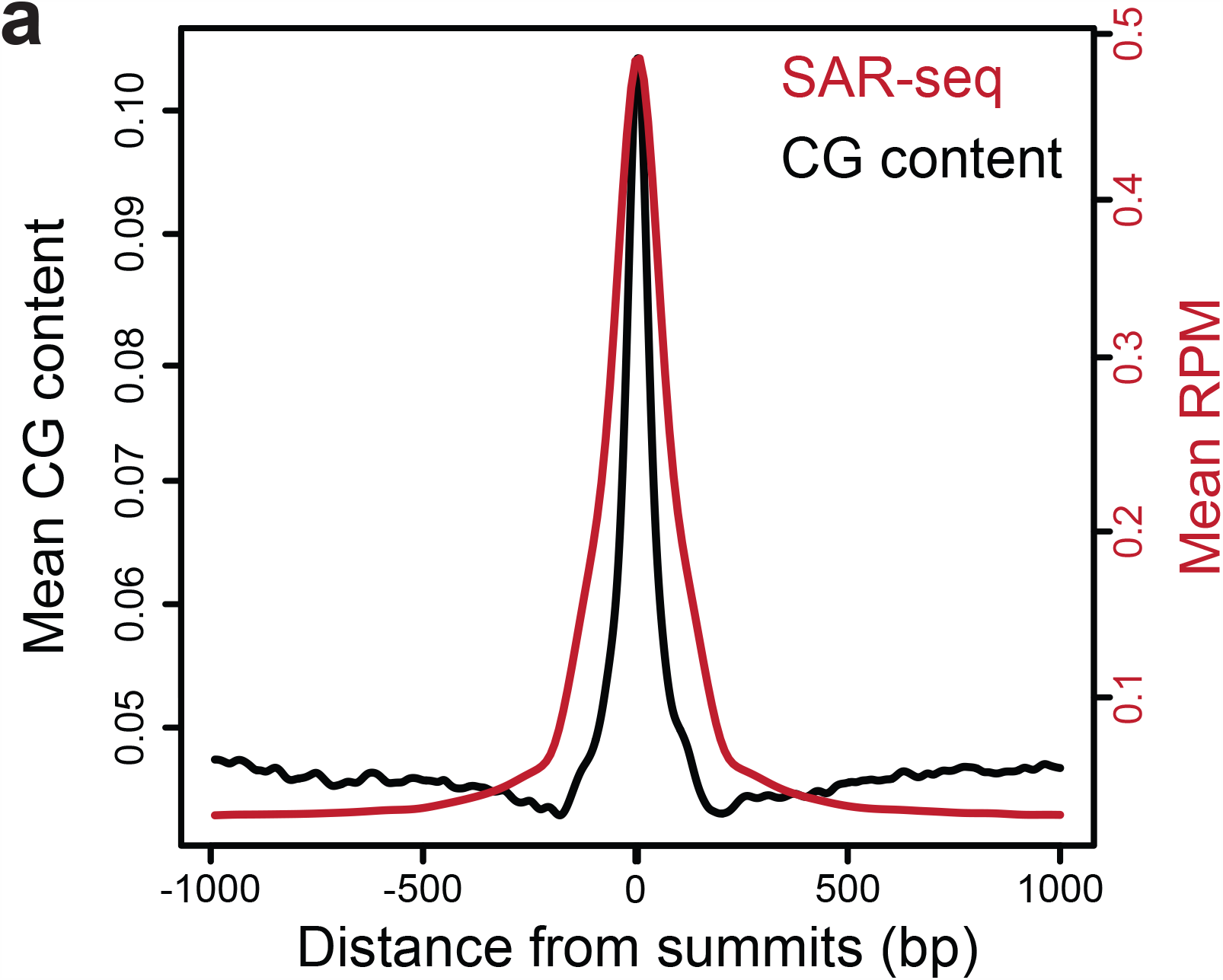
CG dinucleotide content associated with SAR-seq peaks. Aggregate plots showing the distribution of CG dinucleotide (black) at ±1kb around SAR-seq summit overlaid with SAR-seq signal (red).

